# RNA-seq and Bulked Segregant Analysis of a Gene Related to High Growth in *Ginkgo biloba* Half-siblings

**DOI:** 10.1101/034272

**Authors:** Haixia Tang, Jihong Li, Shiyan Xing, Shuhui Du, Zhongtang Wang, Limin Sun, Xiaojing Liu

## Abstract

The lifetime of *G. biloba* is very long, and its growth is relatively slow. However, little is known about growth-related genes in this species. We combined mRNA sequencing (RNA-Seq) with bulked segregant analysis (BSA) to fine map significant agronomic trait genes by developing polymorphism molecular markers at the transcriptome level. RNA-Seq data provides BSA with genotype information in RNA Pool to screen out linked genes (low in false positives) after data analysis, and the efficiency of development and verification of the linked polymorphism marker is greatly improved. This combined approach (named BSR) has been applied to plant transcriptome sequencing in sunflower, corn, wheat, and *Arabidopsis thaliana*. In this study, transcriptome sequencing of high growth (GD) and low growth (BD) samples of *G. biloba* half-sib families was performed. After assembling the clean reads, 601 differential expression genes were detected and 513 of them were assigned functional annotations. Single nucleotide polymorphism (SNP) analysis identified SNPs associated with 119 genes in the GD and BD groups; 58 of these genes were annotated. This study provides molecular level data that could be used for seed selection of high growth *G.biloba* half-sib families for future breeding programs.

## Introduction

*Ginkgo biloba* is a deciduous tree in the family Ginkgoaceae. It is the only species in China to survive the quaternary glacier and, as such, is recognized as a “living fossil” (Jacobs and Browner 2000). *G. biloba* has a very long lifetime, the leaf is fan-shaped, the tree is tall and straight, and its tolerance to drought and barren conditions has made it a significant ornamental, greening, edible, medicinal (Kato-Noguchi *et al*. 2013), and timber tree. In China, the cultivated area of *G.biloba* is more than 200000 hm^2^ and the number of trees has been estimated as 913000 (Xing 2014). *G. biloba* growth is relatively slow with the average increment of timber volume reaching its maximum at about 40 years (Yuan *et al*. 2002; Cao 2007). The tree is generally harvested for maximum timber volume at about 60 years (xing *et al*. 1993). Until now, most studies have focused on the physiology (Newcomer 1953; Echenard *et al*. 2008), phylogeny, (Zhang *et al*. 2015; Guo and Chen 2005), and sex-determining mechanism (Liao *et al*. 2009), and molecular biology studies about the growth mechanism of *G. biloba* are relatively few. The genome sequence of *G. biloba* is still unavailable; therefore, genomics studies are relatively difficult. mRNA sequencing (RNA-Seq) is a next-generation sequencing technology (Cloonan *et al*. 2008; Fu *et al*. 2009; Tang *et al*. 2009; Wilhelm and Landry 2009) that has been used widely to authenticate and quantify normal and rare transcripts, and to provide transcript sequential structure information of specific samples (Liu 2010; Maher *et al*. 2009) in species without a reference genome. The recent application of RNA-Seq to *G. biloba* aseptic seedlings identified a gene that encoded chalcone isomerase (GbCHI1), one of the key enzymes in the flavonoid biosynthesis pathway, that exhibited differences in the protein sequence compared with a previously identified GbCHI(Han *et al*. 2015). Transcriptome sequencing of *G. biloba* kernels revealed 66 unigenes that were found to be responsible for terpenoid backbone biosynthesis (He *et al*. 2015). In addition, *G. biloba* genes associated with the biosynthesis of bilobalide and paclitaxel were found by transcriptome sequencing (Zhang *et al*. 2013). Transcriptome sequencing of the epiphyllous ovules of *G. biloba* var. *epiphylla* identified snRNA genes associated with the adjustment and control of ovular development (Zhang *et al*. 2015). However, no studies into high growth-related genes in *G. biloba* have been reported so far. The growth of *G. biloba* can be affected by a combination of the environment, inheritance, and other factors (Ge *et al*. 2003; Zhang *et al*. 2001); therefore, we aimed to study growth-related genes in a large group of *G. biloba* plants to obtain a comprehensive overview of the genes involved.

We combined RNA-seq with bulked segregant analysis (BSA) to fine mapping genes associated with significant agronomic traits gene at the transcriptome level. BSA has been used to rapidly identify genetic markers linked to a genomic region associated with a selected phenotype (Michelmore *et al*. 1991; Maren *et al*. 2013). The fundamental principle of BSA is that extreme differences of individual phenotype or genotype can be used as the basis on which individuals are selected to obtain a DNA mixture, so that two DNA pools equivalent to near-isogenic line can be built. BSA can be used for efficient marker enrichment in a target region (Bauer *et al*. 1997). BSA has been used for a wide range of plant genomic applications, such as genome sequencing in barley (Steuernagel *et al*. 2009; Mackay and Caligari 2000), Arabidopsis (Wolyn *et al*. 2004), rice (Duan *et al*. 2003), corn (Tang *et al*. 2014), and sunflower (Maren *et al*. 2013). In the combined technology (here named BSR), RNA-Seq is used to provide BSA with genotype data in the RNA pools. Linked genes (low in false positives) can be screened out after data analysis, which greatly improves the efficiency of development and verification of linked polymorphism markers. For species with no reference genome, the RNA-Seq data are assembled to obtain unigenes that are subjected to a series of bioinformatics analysis, including genetic structure annotation, gene expression analysis, and gene function annotation.

*G. biloba* half-sib families from a nursery stock at the seedling stage were used in this study. High growth (GD) and low growth (BD) RNA pools were built from the group level, and BSR was used to identify candidate genes related to the high growth trait. These data will expand the existing transcriptome resources of *G. biloba*, and provide a valuable platform for further studies on developmental and metabolic mechanisms in this species. The information can also be used for functional gene studies and molecular breeding programs.

## Results

### Growth analysis and sample collection of *G. biloba* half-sib families

The average seedling height of the GD group was more than the average height of the two groups for 2 consecutive years, while the average seedling height of the BD group was lower than the average height for 2 consecutive years (Fig. 1B). The photosynthetic rate (Pn), which reflects the speed of carbon dioxide fixation during photosynthesis, is shown in (Fig. 1C). The net Pn in the GD group (8.3 μmol m^−2^ s^−1^) was more than the Pn (4.23 μmol m^−2^ s^−1^) in the BD group. The average chlorophyll content, which reflects photosynthetic capacity, was higher in the GD group (8.6 mgg^−1^FW) than in the BD group (4.1 mgg^−1^FW) (Fig. 1D).

**Fig. 1.**
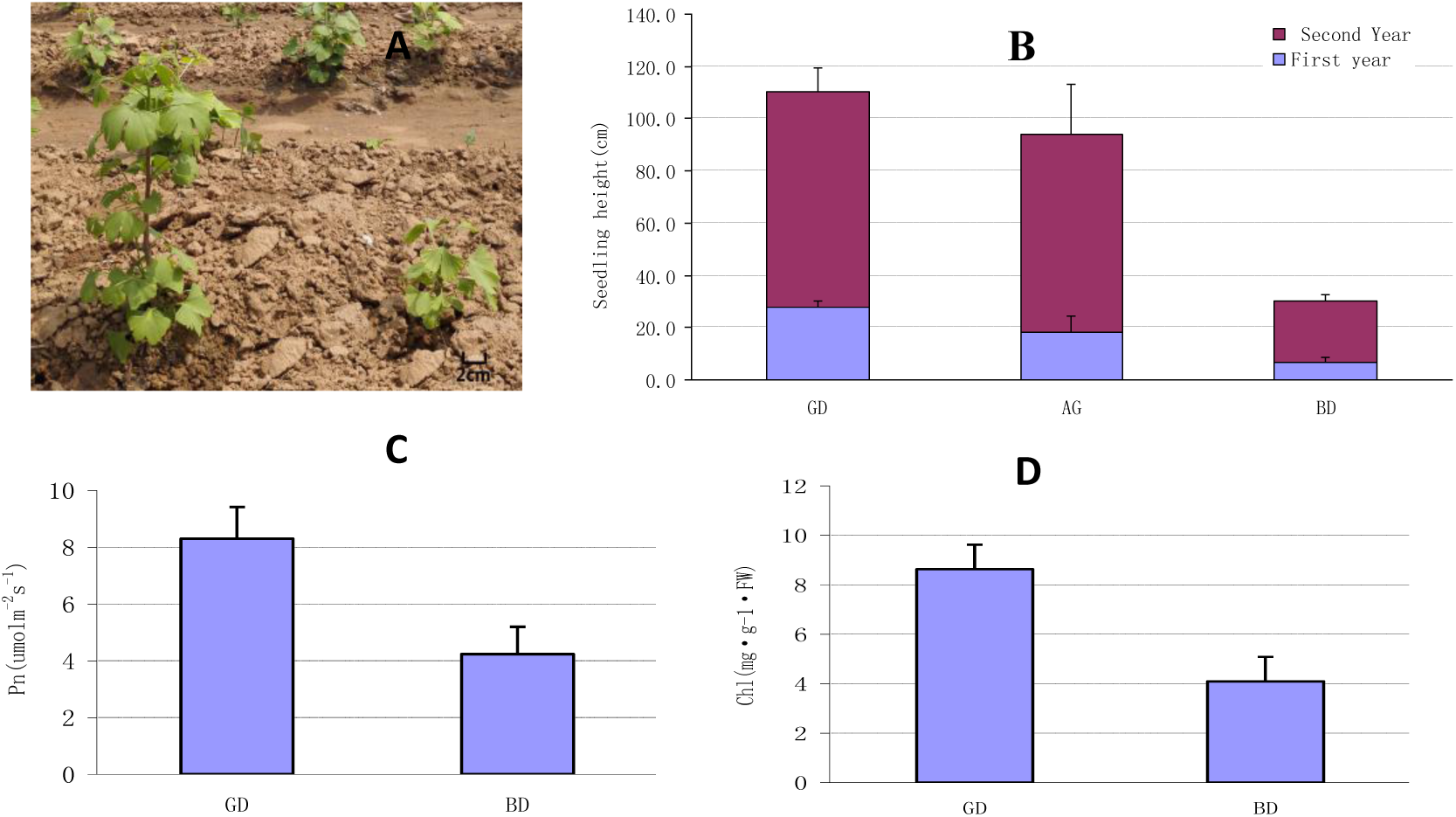
Growth traits of the high growth (GD) and low growth (BD) groups in the *G. biloba* half-sib families. (A) Seedlings in the GD and BD groups. (B) Average height of the seedlings in the GD and BD groups. The average seeding height of the GD group was 27.82 cm in the first year with a net increase of 82.37 cm in the second year; the average seeding height of the BD group was 6.63 cm in the first year with a net increase of 23.43 cm in the second year; AG is the average height of the two groups. (C) Net photosynthetic rate (Pn) in the GD and BD groups. (D) Average chlorophyll content in the GD and BD groups.

### Illumina HiSeq mRNA sequencing

After quality control of the RNA-Seq reads, we obtained 30 Gb of clean reads from the GD and BD groups; the Q30 basic group ratios were more that 90% (Table 1). The clean reads were assembled using Trinity software (Grabherr *et al*. 2011), and a total of 180402 single transcripts and 142492 unigenes are obtained. Of these, 77069 unigenes (27.11% of the total number) were 300–500 bp long, and 18.15% and 15.16% were 500–1000 bp and 1000–2000 bp long, respectively. The N50 of single transcripts was 1514 bp and the N50 of the unigenes was 1081 bp, indicating that the integrity of the assembly was reasonably high (Fig. 2).

**Table 1.** Statistics of the *G. biloba* half-sib families transcriptome sequencing data

| Samples | Clean reads<br>(bp) | Clean data<br>(bases) | GC content | Percent ≥Q30 |
| --- | --- | --- | --- | --- |
| GD | 121909823 | 30477455750 | 44.93% | 92.16% |
| BD | 122947833 | 30736958250 | 45.15% | 91.95% |

Samples: GD, high growth group, BD, low growth group. Clean reads: total number of pair-end reads in the clean data. Clean data: total number of bases in the clean data. GC content: GC content of the clean reads. Percent ≥Q30: percentage of clean data with a quality score greater than or equal to 30 (i.e., the probability that base is called incorrectly is 1 in 1000).

The assembled unigenes were annotated using Clusters of Orthologous Groups (COG) (Tatusov *et al*. 2000), Eukaryotic Orthologous Groups (KOG) (Koonin *et al*. 2004), protein family (Pfam) (Finn *et al*. 2013), Gene Ontology (GO) (Ashburner *et al*. 2000), Kyoto Encyclopedia of Genes and Genomes (KEGG) (Kanehisa *et al*. 2004), SwissProt protein sequence (SwissProt) (Apweiler *et al* 2004), and the NCBI non-redundant protein sequence (Nr) (Deng *et al*. 2006) and nucleotide sequence (Nt) (http://blast.ncbi.nlm.nih.gov/Blast.cgi) databases. The annotation statistics are listed in Table 2. The E-value for the searches against each of the databases was set as ≤1e−5. A total of 41758 (29.3%) unigenes were annotated in at least one of the databases; the remaining 137734 unigenes (60.7%) were not annotated, indicating that *G. biloba* genetic information is deficient in the existing databases. The Nr database produced the highest number of annotated unigenes (38991), while KEGG produced the least (5821).

**Fig. 2.**
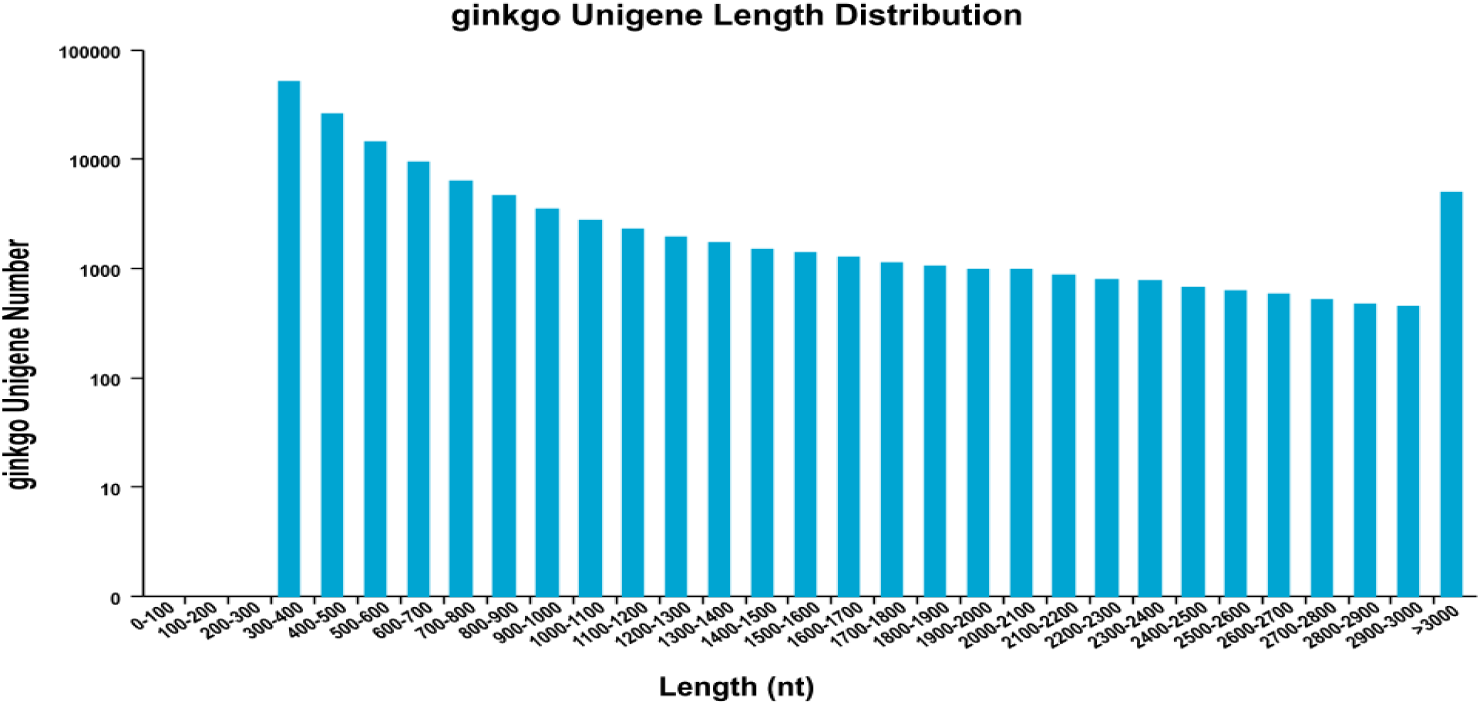
Length distribution of *G. biloba* half-sib families unigenes in the high growth (GD) and low growth (BD) transcriptomes.

**Table 2.** Annotation statistics of the *G. biloba* half-sib families unigene

| Databases | Unigenes | $\geq 300$ bp<sup>a</sup> | $\geq 1000$ bp<sup>b</sup> |
| --- | --- | --- | --- |
| COG | 11719 | 4489 | 7230 |
| KOG | 24483 | 13310 | 11173 |
| Pfam | 21149 | 6775 | 14374 |
| GO | 20002 | 8987 | 11015 |
| KEGG | 5821 | 1755 | 4066 |
| SwissProt | 22705 | 9654 | 13051 |
| TrEMBL | 38598 | 20023 | 18575 |
| Nr | 38991 | 20416 | 18575 |
| Nt | 22101 | 7394 | 14707 |

<sup>a</sup>≥300 bp indicates the number of annotated unigenes ≥300 bp long. <sup>b</sup>≥1000 bp indicates the number of annotated unigenes ≥1000 bp long.

### Expression analysis of differentially expressed genes of the *G. biloba* half-sib families

False discovery rate (FDR) values were adopted as a key index of differentially expressed genes (DEGs) to reduce false positives that may be caused by independent statistical hypothesis testing of expression values of a large number of genes. FDR values <0.05 and the differential multiple fold changes (FC) ≥2 between two groups were used as the cutoff to identify DEGs. A scatter plot of gene expression levels in the GD and BD groups shows that most of the points fell on the diagonal (Fig. 3A), indicating that the gene expression trends were similar in the two groups and the repetition correlation was high. A volcano plot of the differential gene expression between the BD and GD groups (Fig. 3B) shows that the number of genes with significant −log FDR and FC values was more than the number of genes without significant −log FDR and FC values, indicating that the screening was reliable. The A total of 601 DEGs were identified between the BD (control) and GD (test) groups; 400 were up-regulated and 201 were down-regulated. Hierarchical clustering analysis of the DEGs showed that genes with the same or similar expression patterns clustered together (Fig. 3C).

**Fig. 3.**
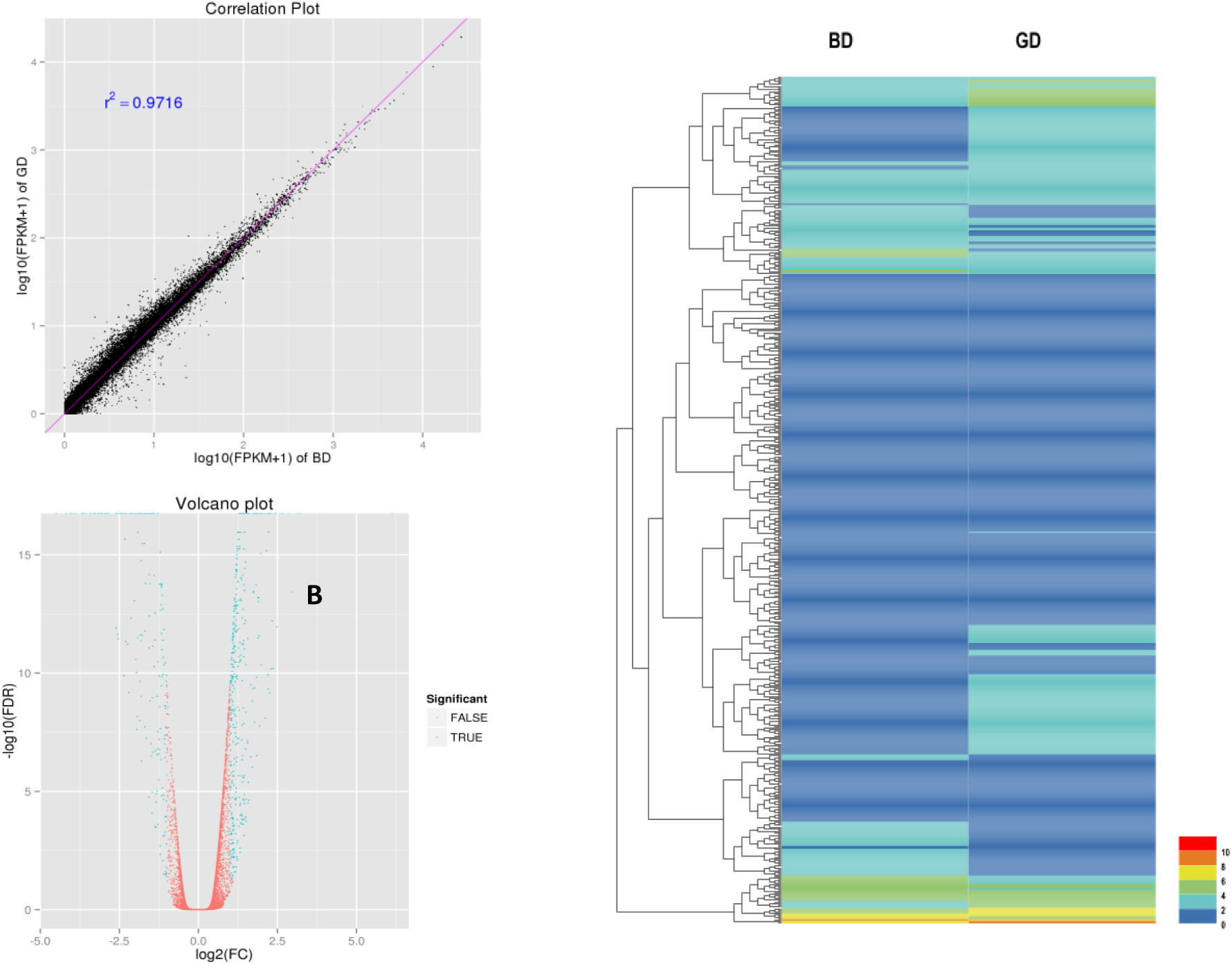
Analysis of gene expression in the high growth (GD) and low growth (BD) transcriptomes. (A) Scatter diagram of gene expression levels in the GD and BD groups. FPKM, fragments per kilobase of transcript per million mapped reads. (B) Volcano plot of differential gene expression between the GD and BD groups. Green indicates genes with significant −log FDR and FC values; red indicates genes without significant −log FDR and FC values. FDR, false discovery rate; FC, fold change. (C) Hierarchical clustering of DEGs with the same or similar expression patterns between the BD (control) and GD (test) groups.

### Functional annotation of differentially expressed genes of *G. biloba* half-sib families

The second-level GO functional annotation terms assigned to the DEGs and to all the unigenes are shown in Fig. 4. Differences between the percentages of genes assigned to the different functions may be related to high growth. Under cellular component, “cell” (73, 6.3%), “cell part” (75, 6.4%), and “organelle” (55, 4.7%) were assigned to the highest number of genes; under molecular function, “catalytic activity” (124, 10.6%) and “binding” (124, 10.6%) were assigned to the highest number of genes; and under biological process, “metabolic process” (169, 14.5%), “cellular process”(126, 10.8%), and “single-organism process” (91, 7.8%) were assigned to the highest number of genes. Among the 25 COG categories (Fig. 5), “Replication, recombination and repair” (39, 22.9%) and “General function prediction only” (39, 22.9%) were assigned to the highest number of DEGs, followed by “Posttranslational modification, protein turnover, chaperones” (29, 17.1%) and “Transcription” (28, 16.5%). The categories with the lowest number of DEGs were “Coenzyme transport and metabolism” (1, 0.59%), “Inorganic ion transport and metabolism” (2, 1.2%), and “Chromatin structure and dynamics” (2, 1.2%). None of the DEGs were assigned to “Nuclear structure”, “Defense mechanisms”, “Intracellular trafficking, secretion, and vesicular transport”, or “Nucleotide transport and metabolism”.

To explore the biological pathways in which the DEGs may be involved, we performed a KEGG analysis (Fig. 6). Many DEGs were assigned to the Spliceosome, Protein processing in endoplasmic reticulum, RNA transport, and Ubiquitin-mediated proteolysis pathways. Splicing factors Prp22, Sm, SF3a, Prp6, P68, S164, Snu66, CDC5, and THOC are known to participate in mRNA splicing and genes encoding them were among the up-regulated genes in the GD group compared with BD group. The protein responsible for endoplasmic reticulum-associated degradation (ERAD) is related to *Hsp70* and *sHSF*, which were down-regulated in GD compared with BD. Meanwhile, the genes encoding the ubiquitin-conjugating E2 enzyme (UBE20) and the ubiquitin E3 ligase (ARF-BR1) associated with the proteasome were up-regulated in GD compared with BD. Genes encoding the THOC2, Tpr, Nup62, eIF5B, and eIF4G factors, which are involved in RNA transport, were up-regulated in GD compared with BD.

**Fig. 4.**
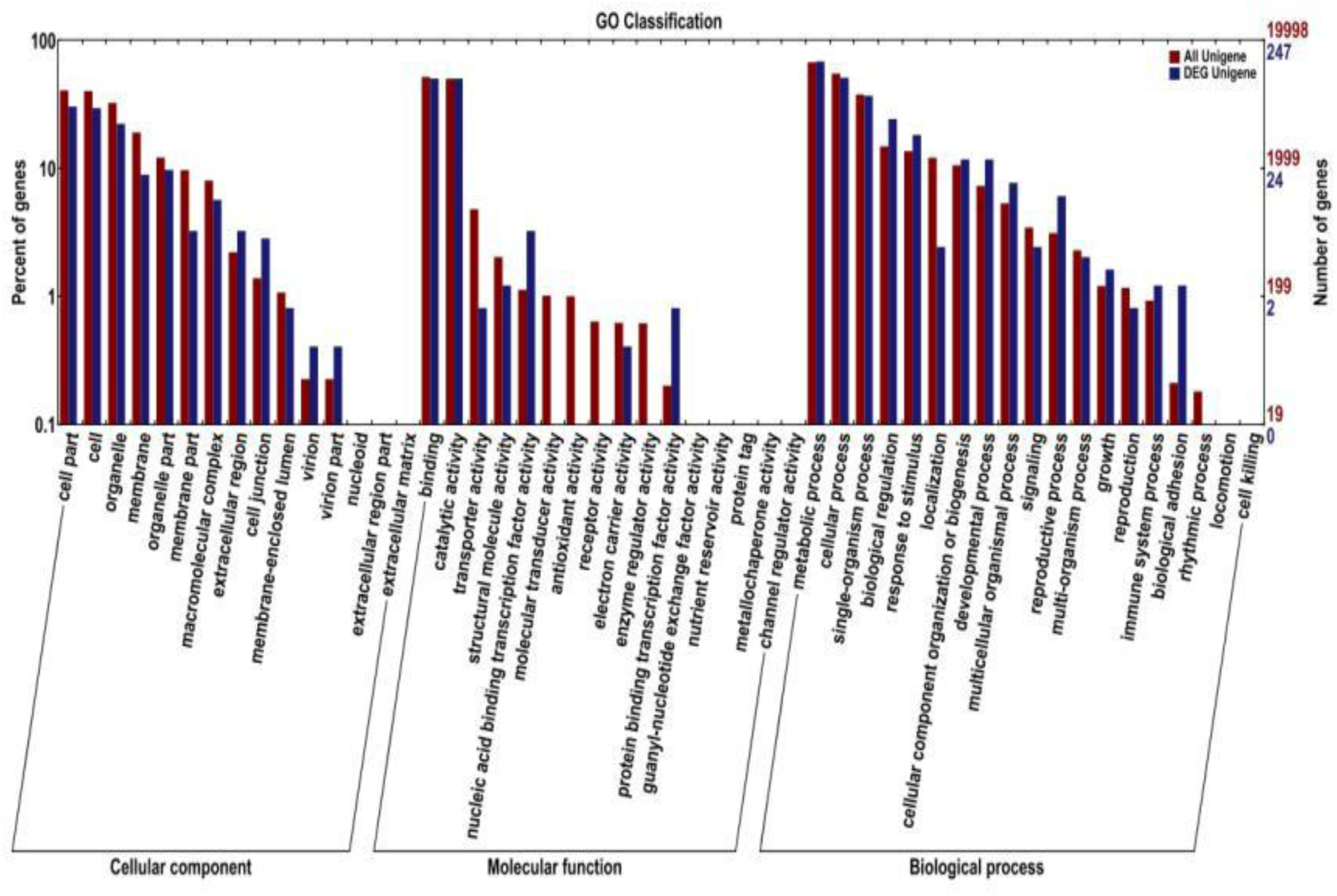
Gene Ontology (GO) terms assigned to differentially expressed genes and all unigenes in the *G. biloba* half-sib families transcriptomes. Second-level terms were assigned under the three GO categories: cellular component, molecular function, and biological process.

**Fig. 5.**
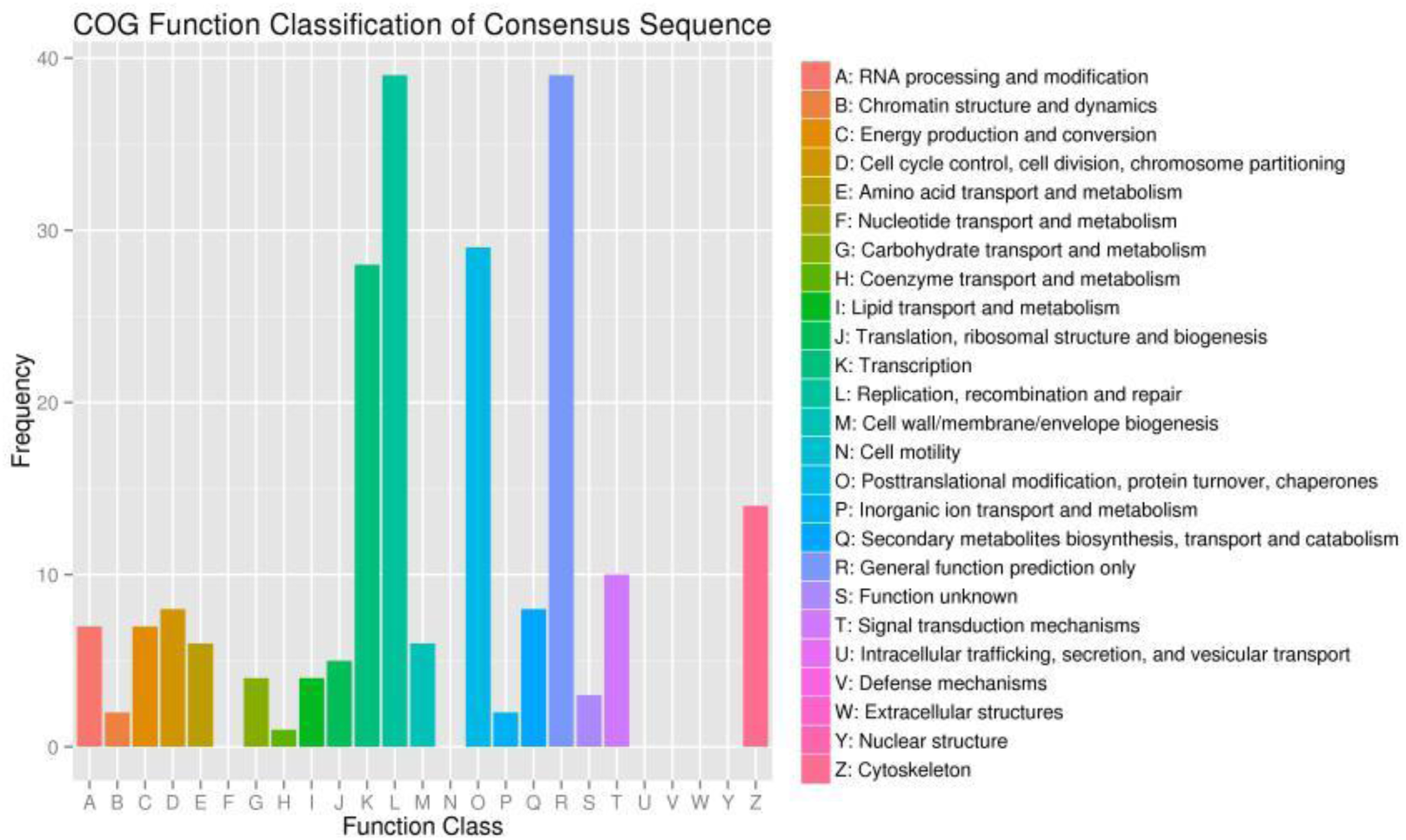
COG annotations assigned to differentially expressed genes in the *G. biloba* half-sib families transcriptomes.

**Fig. 6.**
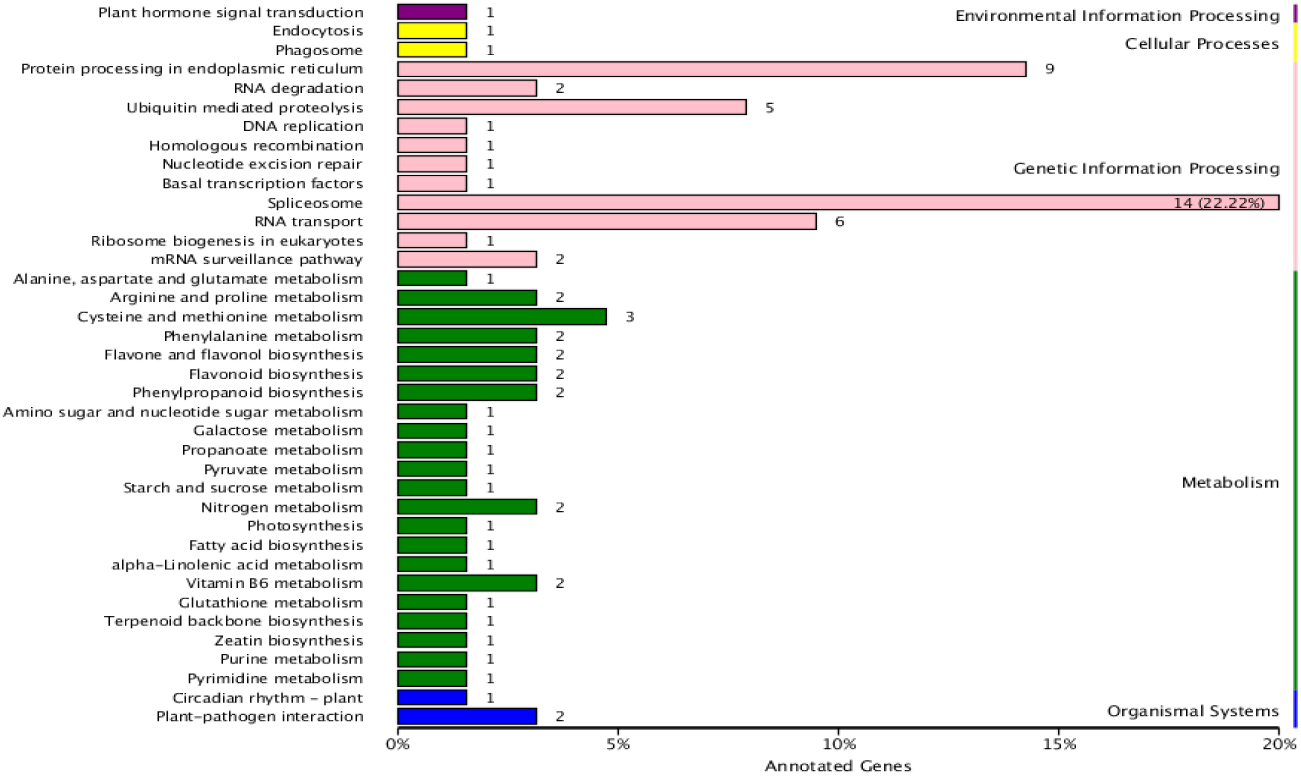
KEGG annotations of differentially expressed genes in the *G. biloba* half-sib families transcriptomes.

### Relevance of the predicted SNPs of *G.biloba* half-sib families

The unigene sequences were compared with the known sequences in three protein sequence databases (Nr, SwissProt, and KEGG) and the protein-coding sequence (CDS) information from the matched sequences was used to annotate the unigenes. The CDSs were translated into amino acid sequences according to the standard codon table. The CDSs of unigenes that did not match any of the known protein sequences were predicted using the GetORF software (http://embossgui.sourceforge.net/demo/getorf.html), which translates nucleotide sequences in all six reading frames. The longest amino acid sequence for each unigene was selected as the translated protein sequence for that gene. The length distributions of the CDSs and predicted protein sequences of all the unigenes are plotted in Fig. 7.

**Fig. 7.**
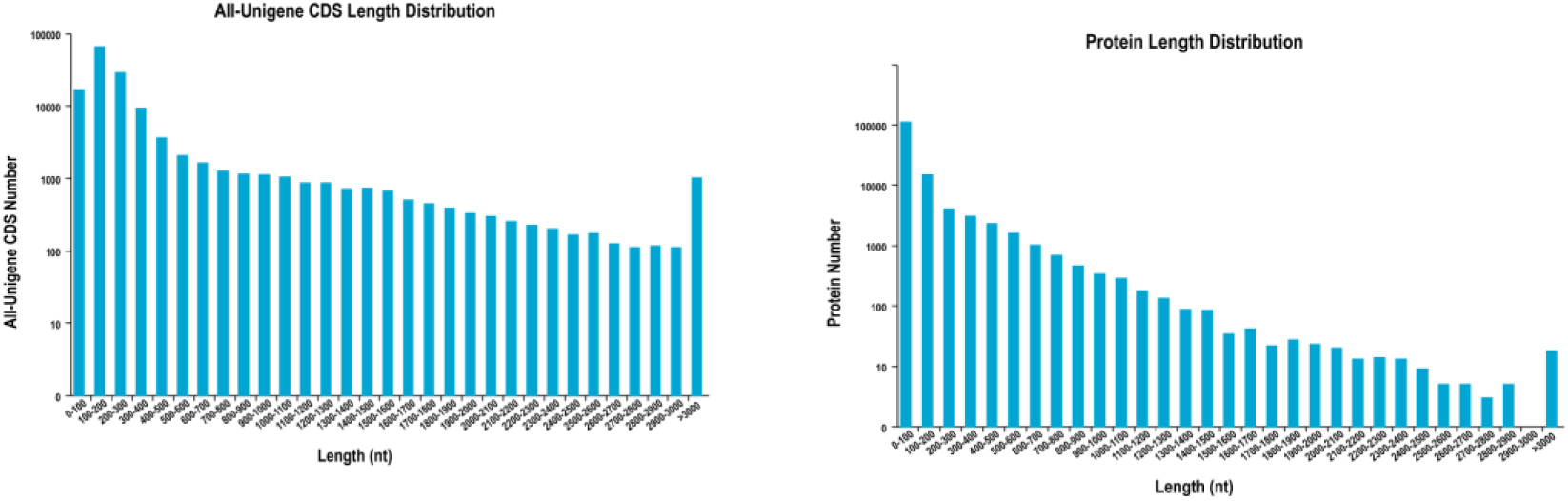
Length distribution of the protein-coding sequences (CDSs) and translated amino acid sequences of all the unigenes in the *G. biloba* half-sib families transcriptomes.

The RNA-Seq reads from each group were compared with the assembled unigenes, to identify candidate single nucleotide polymorphisms (SNPs). A total of 115517 SNP loci were acquired. After filtering SNPs with a depth of less than 3 (21776) and undifferentiated loci (67859) between the CD and BD pools, 25883 SNP loci remained. SNP loci with ED^5^ values (Euclidean distance) higher than the threshold value (set as 1.151) were regarded as outstanding correlative loci (Table S1). The number of genes associated with these SNPs was 119, of which 58 were annotated genes (Table S2). Of the 58 annotated genes, 31 had KOG annotations only under General function, Carbohydrate transport and metabolism, and Posttranslational modification, protein turnover and chaperones. Twenty-nine of the 58 genes were assigned GO terms. Under biological process, metabolic process (GO: 0008152), regulation of transcription, DNA-templated (GO: 0006355), and regulation of plant-type hypersensitive response (GO: 0010363) were highly represented; under cellular component, plasma membrane (GO: 0005886) and integral component of membrane (GO: 0016021) were highly represented; and under molecular function, binding (GO: 0005488) and metal ion binding (GO: 0046872) were highly represented (Table S3). The 58 genes were annotated with seven KEGG pathways, together with Protein processing in endoplasmic reticulum and Spliceosome, which were annotated to DEGs, Sphingolipid metabolism, Alanine, aspartate and glutamate metabolism and Carbon sequestration in photosynthetic organisms were also represented.

### Expression of high growth trait-related genes

DEGs related to high growth in *G. biloba* half-sib families were predicted to participate in photosynthetic carbon sequestration. After photosynthetic carbon sequestration of CO_2_, reactive enzyme activation occurs through the glycolysis process. The correlation gene *(c28693_g1_i1)* has a regulatory effect on the dehydrogenation and phosphorylation of 1,3-2-glyceric-acid phosphate to form glyceraldehyde 3-phosphate. This gene also participates in oxidoreductase activity, acting on the aldehyde or oxo group of donors, NAD or NADP as acceptor, and NADP binding activities. It has been shown that improvement of plasmalemma redox activity can promote elongation growth of plants (Qui *et al*. 1985; Cao *et al*. 1997). The Pn and growth rate of group GD were higher than those of group BD, which may be related to the activation of genes involved in the photosynthetic carbon sequestration process of *G. biloba*.

Sphingolipids play major roles in intracellular transduction (Merrill 2002) and participate in many important signal transduction processes, such as adjustment of cellular growth, differentiation, senility, and programmed cell death (Liu and Gou 2009). Sphingolipid metabolism can be controlled by differential enzyme expression and is cell specific expression, and ceramidase has been implicated in different tissues (Riebding *et al*. 2003). Ceramidase activity has been correlated with high growth, which indicates that sphingolipids in the GD group may be related to high growth. In addition, a related enzyme involved in the activity of splicing factor Prp22 and a correlation factor associated with the snRNA component were both up-regulated in the GD group. We speculate that the spliceosome-encoding gene may effectively promote high growth in *G. biloba*.

Endoplasmic reticulum-associated protein degradation eliminates denatured proteins, paraproteins or damaged proteins, plays a major role in controlling the quality of proteins. The KEGG pathway analysis revealed that ERAD was related to the down-regulated DEGs *Hsp70* and *sHSF*. It has been shown that degradation of ERAD substrate was coupled with the degradation pathway of ubiquitin-proteasomes (Hiller *et al*. 1996). The DEGs *E2 (UBE20)* and *E3 (ARF-BR1)* proteasome that participate in ubiquitin-mediated proteolysis were up-regulated in the GD group. The ERAD system can preferentially degrade specific proteins and effectively protect the immune system, suggesting that it may be related to the high growth of the *G.biloba* seedlings.

### Discussion

For the BSA, the ED value of each SNP was calculated between the GD and PD RNA pools using the allele depth of the differentially occurring SNP, determine the target site, and conduct linked marker. A total of 119 genes were correlated with the identified SNPs, and 58 of them were assigned functional annotations. In Bulked Segregant Analysis and Amplified Fragment Length Polymorphism (BSA-AFLP) analysis of the resistance gene *rhm* of corn southern leaf blight, more than 222 polymorphic markers were found in a F1-generation infection resistance pool (10 plants in each pool); however, further verification found that in 80 single plants of the F2-generation, 16 of the markers were not linked with the target gene (Cai *et al*. 2003). A similar result has been reported in barley (Molna *et al*. 2000). It indicates that the non-linked marker can also present to polymorphic stripe of two pools. Although these issues cannot be entirely eliminated, they can be reduced by increasing the number of single plants in the mixing pools. In the present study, 30 single plants were used in each mixing pool, which made up 30.9% of the total samples and reduced the number of non-linked markers that were identified. In addition, to ensure the veracity of the gene screening, expression analysis and identification of SNP loci were performed using the RNA-Seq data to detect growth-related genes and lay the foundation for fine mapping of these genes in the *G. biloba* half-sib families.

In most woody plants, heterozygosity is strong and the genomes tend to be large and complex; therefore, studies into the genetic background of these plants have been limited. For species without a reference genome, RNA-Seq data have been used to obtain inheritance information and to build physical maps (Li *et al*. 2010). *G. biloba* is an ancient gymnosperm that is widely distributed around the world and its ability to growth and adapt to different environmental conditions suggests that a large number of responsive genes would have evolved (Li 2011). Many genes and transcription factors related to growth and development of *G. biloba* are available in the related study of *G. biloba* leaves, for example, *COP9* signal corpuscle composite subunits, *AGAMOUS-like MADS-box* transcription factor (Shore and Sharrocks 1995), *glucan endo-l,3-beta-glucosidase* (Meirinho *et al*. 2010), *DELLA, ELFB, homeobox-leucine zipper protein*, and *EMBRYONICFLOWER 2* (Lin *et al*. 2010). Based on the expression levels of genes in different samples, 601 DEGs have been recognized and functional annotations have been assigned to 513 of them. Among them, two *Homeobox-leucine zipper protein genes* were up-regulated in the GD group compared with the BD group; therefore, these are very likely related to high growth of *G. biloba*. In addition, the DEGs and the gene associated with BSR technology were found to be associated with spliceosome activity, spliceosome metabolism, photosynthetic carbon sequestration, and endoplasmic reticulum protein processing and also to participate in growth and metabolism of *G. biloba*.

## Materials and Methods

### Genetic materials

*G. biloba* seeds were obtained from the *G. biloba* germplasm resource garden of the Gaoqiao Tree Farm in Tai’an City, Shandong Province, China. The experimental field is located N 35°54′, E 116°53′, which has a continental warm temperature zone medium-latitude monsoon climate. The average annual temperature is 13.4°C, and the maximum and minimum recorded air temperatures are 40.7°C, and −19°C, respectively. The annual average rainfall is 689.6 mm, average annual evaporation is 1169.8 mm, and the average number of frost-free days is 206 per year. The soil is sand loamy river moisture soil. A total of 358 seeds were collected in Shiqiao Town, Pan County, Guizhou Province on 29 September 2013. Seeding was conducted in 2014 and 194 seedlings emerged. After planting, field management measures were uniform throughout. Seedling height was measured in December 2014 and November 2015. The 30 tallest seedlings and 30 shortest seedlings were selected to form the GD and BD groups, respectively. The heights of the selected seedlings were recorded for 2 consecutive years. The number of seedlings in the half-sib families group was 194, and the variable coefficient of seedling height in the families was >30%. The initial expanded second lamina at the top of the seedlings in group GD and group BD were punched and then disposed in mixing pool mode in May 2015, then marked as GD or BD, quick-frozen in liquid nitrogen, and stored at −80°C until used.

### Extraction of RNA from *G. biloba* half-sib families leaf tablets

Total RNA from each sample was isolated separately using a RN38 EASY spin plus Plant RNA kit (Aidlab Biotech, Beijing, China). Nanodrop Analyzer (Thermo Science, Wilmington, USA), Qubit 2.0 Fluorometer and Agilent 2100 Bioanalyzer (Agilent Technologies, Santa Clara, CA, USA) were used to estimate the purity, concentration, and integrity of the extracted RNA.

### cDNA library construction and sequencing

Total mRNA was isolated by oligo (dT) selection using Dynabeads mRNA DIRECT Kit (invitrogen), and each sample was prepared 5 ug for constructing the cDNA library. The purified mRNA was fragmented at elevated temperature (90°C), then reverse transcribed to first strand cDNA with random primer. Second strand cDNA was synthesized in the presence of DNA polymerase I and RNaseH. The cDNA was cleaned using Agencourt Ampure XP SPRI beads (Beckman Coulter). The cDNA molecules were subjected to end repair, and add an ‘A’ base at the 3’-end. Illumina adapters were ligated to the cDNA molecules, resultant cDNA library was amplified using PCR for enrichment of adapter ligated fragments. Libraries were prepared from a 400–500 bp size-selected fraction following adapter ligation and 2% agarose gel separation. The cDNA library was quantified using qPCR method(>10 nM). It was then sequenced using the Illumina Hi-Seq2500 platform.

### Unigene function annotation

The raw reads were cleaned by removing adapter sequences, reads containing ploy-N, and low-quality sequences (Q <30). Clean reads were aligned to the reference genome sequence using the program Tophat(Yang *et al*. 2015; Rong *et al*. 2015).

The assembled unigene sequences were searched against the Nr, SwissProt, GO, COG, KOG, Pfam, and KEGG databases using the NCBI Basic Local Alignment Search Tool (BLAST) tools (Altschul *et al*. 1997) to annotate the unigenes.

### Unigene structural analysis

The CDSs of the unigenes were predicted based on their alignment to known protein sequences. The predicted CDSs were translated into amino acid sequence using the standard codon table. The unassembled clean reads in each sample were mapped to the assembled unigene sequences. SNP loci were detected using the SNP calling program in the Genome Analysis Toolkit (GATK) (https://www.broadinstitute.org/gatk/index.php). SNP loci were screened then we chose to measure allele segregation using Euclidean distance (ED), as a metric that does not require parental strain in-formation and is resistant to noise (Jonathon T. et al. 2013). In order to obtain good correlation effect, The ED value was disposed in the 5 power mode, and the data were recognized as the basis for BSR relevance.Using the equation:

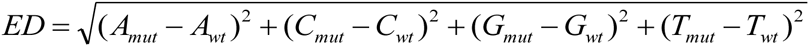
where each letter (A, C, G, T) corresponds to the frequency of its corresponding DNA nucleotide.

### Analysis of differential gene expression

Reads is compared with Unigene bank obtained by sequencing of each sample using Bowtie software (Langmead *et al*. 2009). The expression levels were estimated by combining with RSEM (Li and Colin 2011). RSEM (RNA-Seq by Expectation Maximization), which implements our quantification method and provides extensions to our original model. The expression levels of the unigenes were expressed as fragments per kilobase of transcript per million mapped reads (FPKM) values to eliminate the influences of gene length and sequencing quantity difference on of the estimate gene expression. FPKM values can be used directly to compare gene expression differences between samples.

FPKM was calculated as follows:

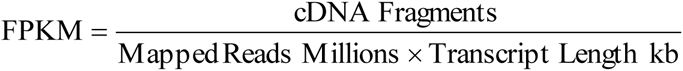
where “cDNA Fragments” is the number of fragments of one transcript in the sample (i.e., the number of double-end reads); “Mapped Reads Millions” is the number of mapped reads (in this study it was 106); and “Transcript Length kb” is the length of the transcript.

Differential expression analysis between the GD and BD groups was conducted using DESeq (Anders and Huber 2010). Significance p-values were obtained by original hypothesis testing and adjusted using the Benjamini–Hochberg method. The FDR was used as the key index for screening the DEGs, and the screened DEGs were analyzed in a hierarchical clustering mode.

**Data Availability**: The raw reads of the RNA-seq are now beening processed by NCBI staff. File S1 contains SNP depth in the RNA-Seq data of *Ginkgo biloba* half-sib families. File S2 contains functional annotation of unigenes of *Ginkgo biloba* half-sib families. File S3 contains Gene Ontology annotation of unigenes of *Ginkgo biloba* half-sib families.

## Acknowledgments

We thank Rongkai Wang, for constructive criticism and suggestions, Haidong Gao, for experimental operation and the data analysis, and Shuhui Du, for the Paper modified. This work was supported by National Science and Technology Support Plan (2012BAD21B04), Shandong Agricultural Seeds Engineering Project: LuNong good word [2011] no. 7, National Plant Germplasm Resources Sharing Platform - National Forest Tree Germplasm Resources (including bamboo rattan flower) Platform (platform subsystem): 2013–39.

## Author Contributions

Conceived and designed the experiments: SYX, JHL and HXT. Performed the experiments: HXT and JHL. Analyzed the data:SHD, HXT, ZTW, LMS, XJL. Wrote the paper: HXT, JHL and SYX.

